# Expression and *in vitro* anticancer activity of Lp16-PSP, a member of the YjgF/YER057c/UK114 protein family from the mushroom *Lentinula edodes* C_91-3_

**DOI:** 10.1101/2020.04.27.062232

**Authors:** Thomson Patrick Joseph, Qianqian Zhao, Warren Chanda, Sadia Kanwal, Yukun Fang, MinTao Zhong, Min Huang

**Affiliations:** Department of Microbiology, College of Basic Medical Sciences, Dalian Medical University, Dalian, Liaoning 116044, P.R. China; Center for Neuroscience, Shantou University Medical College, Shantou, P.R. China; Computational System Biology Laboratory, Department of Bioinformatics, Shantou University Medical College, Shantou P.R. China; Department of Biotechnology, College of Basic Medical Sciences, Dalian Medical University, Dalian 116044, Liaoning 116044, P.R. China

**Keywords:** *Lentinula edodes* C_91-3_, latcripin-16, endoribonuclease, HL-60 cell line

## Abstract

Latcripin-16 (Lp16-PSP) is a gene that was extracted as a result of *de novo* characterization of the *Lentinula edodes strain* C_91-3_ transcriptome. The aim of the present study was to clone, express and investigate the selective *in vitro* anticancer potential of Lp16-PSP in human cell lines. Lp16-PSP was analyzed using bioinformatics tools, cloned in a prokaryotic expression vector pET32a (+) and transformed into *E. coli* Rosetta gami. It was expressed and solubilized under optimized conditions. The differential scanning fluorometry (DSF)-guided refolding method was used with modifications to identify the proper refolding conditions for the Lp16-PSP protein. In order to determine the selective anticancer potential of Lp16-PSP, a panel of human cancerous and non-cancerous cell lines was used. Lp16-PSP protein was identified as endoribonuclease L-PSP protein and a member of the highly conserved YjgF/YER057c/UK114 protein superfamily. Lp16-PSP was expressed under optimized conditions (37°C for 4 h following induction with 0.5 mM isopropyl β-D-1-thiogalactopyranoside). Solubilization was achieved with mild solubilization buffer containing 2M urea using the freeze-thaw method. The DSF guided refolding method identified the proper refolding conditions (50 mM Tris-HCl, 100 mM NaCl, 1 mM EDTA, 400 mM Arginine, 0.2 mM GSH and 2 mM GSSG; pH 8.0) for Lp16-PSP, with a melting transition of ~58°C. A final yield of ~16 mg of purified Lp16-PSP from 1 L of culture was obtained following dialysis and concentration by PEG 20,000. A Cell Counting Kit-8 assay revealed the selective cytotoxic effect of Lp16-PSP. The HL-60 cell line was demonstrated to be most sensitive to Lp16-PSP, with an IC_50_ value of 74.4±1.07 μg/ml. The results of the present study suggest that Lp16-PSP may serve as a potential anticancer agent; however, further investigation is required to characterize this anticancer effect and to elucidate the molecular mechanism underlying the action of Lp16-PSP.

## 1.0 Introduction

Cancer is a global health problem with high morbidity and mortality worldwide. In the United States alone, ~1,806,590 new cases and ~606,520 cancer-associated mortalities were expected in 2020 (Siegel et al., 2020). Chemotherapy has been extensively used to treat various cancers; however, chemotherapeutic agents often cause severe side effects and cancers may become chemoresistant (Luqmani, 2005). Therefore, the development of novel agents with no or minimal side effects is important to improve the prognosis of patients with cancer.

In the past decade, the pharmacological effects of a number of natural products and compounds derived from natural products have been investigated in clinical trials, especially as anticancer agents (Harvey, 2008, Dias et al., 2012). A number of mushroom species, both edible and medicinal, have been reported to have antiproliferative, antioxidant, cytotoxic, anti-diabetic, anti-microbial, anti-inflammatory and immunomodulatory potentials (Kim et al., 2004, Dong et al., 2007, Liu et al., 2009, Jiang and Sliva, 2010, Thohinung et al., 2010, Bassil et al., 2012). Several components of mushrooms, including polysaccharides, polysaccharide-protein complexes, dietary fibers, certain proteins, terpenoids, steroids and phenols have been reported to have anticancer effects (Ivanova et al., 2014, Singh et al., 2016, Joseph et al., 2017). *Lentinula edodes,* or the shiitake mushroom, grows well in warm and moist climates and has been used as a model mushroom for isolating and investigating the functional properties of lead compounds to be used in pharmaceuticals (Wasser, 2005). It has previously been reported that polysaccharides, terpenoids, sterols and lipids from *L. edodes* are effective treatments for a number of human ailments, including cancer (Wasser, 2005, Resurreccion et al., 2016). Various expression systems have been used for cloning and expressing the enzymes and proteins of *L. edodes* (Zhao and Kwan, 1999, Zhao et al., 2000, Sakamoto et al., 2006). In 2012, our research group demonstrated the anticancer potential of mycelia, a protein component of *L. edodes* strain C_91-3_, in *in vitro* and *in vivo* models and used deep solexa sequencing for the *de novo* characterization of the *L. edodes* C_91-3_ transcriptome (Liu et al., 2012, Zhong et al., 2013). A total of 57 million reads were produced and assembled into 18,120 coding unigenes. These coding unigenes were annotated with gene description, gene oncology and a cluster of orthologous groups based on a similarity search of known proteins. Finally, thousands of genes were extracted from *L. edodes* strain C_91-3_ to further investigate their therapeutic applications.

In the present study, the nucleotide and amino acid sequence of Latcripin-16 was analyzed using bioinformatics tools. Lp16-PSP was identified as the endoribonuclease liver perchloric acid-soluble protein (L-PSP) protein and belongs to the highly conserved YjgF/YER057c/UK114 protein family. This protein family was established following the discovery of YjgF from *Escherichia coli,* YER057c from *yeast* and UK114 from goat (Liu et al., 2016). Latcripin-16 was therefore abbreviated to Lp16-PSP based on its family. Homologs of the YjgF/YER057c/UK114 family are widely distributed across all three domains of life, with high sequence and structural similarities and functional diversity (Thakur et al., 2010). YjgF/YER057c/UK114 proteins are small proteins (~15 kDa) that are typically found in eubacteria, archaea and eukaryotes with 6-9 conserved amino acid residues (Oka et al., 1995, Asagi et al., 1998, Goupil-Feuillerat et al., 1997). The first member of this family to be discovered was a rat liver perchloric acid-soluble protein, characterized as an endoribonuclease, which inhibited the initiation of protein translation in rabbit reticulocyte lysate systems by directly affecting the mRNA template activity (Oka et al., 1995, Morishita et al., 1999). The hp14.5, a homologue of rat liver perchloric acid-soluble protein, UK114 from goat liver and a bovine homologue have been identified as translation inhibitors (Manjasetty et al., 2004), antineoplastic and tumor antigens (Ceciliani et al., 1996, A. Bartorelli and M. Bailo, 1994, S. Racca, 1997, Ghezzo et al., 1999) and activators of calpains (Farkas et al., 2004) respectively. A number of other proteins of the same family with diverse functions have also been identified in bacteria and eukaryotes, including the purine regulator YabJ from *Bacillus subtilus* (Sinha et al., 1999b), YIL051c and YER057c from *Saccharomyces cerevisiae* (Kim et al., 2001) and plant proteins that serve a role in photosynthesis/chromoplastogenesis (Leitner-Dagan et al., 2006). Recently, endoribonuclease L-PSP protein from the *Rhodopseudomonas palustris* strain JSC-3b has been reported to have antiviral activities (Su et al., 2015). Additional functions of the YjgF/YER057c/UK114 superfamily have been reported, including the suppression of cell proliferation and fatty acid binding (Kanouchi et al., 2001, Sasagawa et al., 1999).

Based on the bioinformatics analysis of Lp16-PSP and the reported functions of YjgF/YER057c/UK114 family members, it was hypothesized that Lp16-PSP, being an endoribonuclease L-PSP protein, could cause the degradation of RNAs and inhibit protein translation in human cell lines. The aim of the present study was to clone, express and recover the bioactive form of Lp16-PSP protein. The Lp16-PSP gene encoding the endoribonuclease L-PSP domain was cloned into pET32a (+) plasmids, transformed and expressed in *E. coli* Rosetta-gami (DE3). As the majority of the protein was expressed in the form of inclusion bodies, the denaturation of inclusion bodies was systematically investigated and the dilution method was applied for refolding purpose. The selective anticancer activity of Lp16-PSP was investigated using a panel of human cancerous and normal cell lines.

## 2.0 Materials and methods

### 2.1 Bacterial strains, plasmids and reagents

The *Lentinula edodes* strain C_91-3_ was purchased from The Chinese General Microbiological Culture Collection Center (Chinese Academy of Sciences, Beijing, China). The bacterial strains used in this study were JM109 and Rosetta gami DE3, purchased from Takara Bio, Inc. (Otsu, Japan) and Invitrogen (Thermo Fisher Scientific, Inc., Waltham, MA, USA), respectively. The cloning vector pMD20-T was purchased from Takara Bio, Inc. and the expression vector pET32a (+) was purchased from Invitrogen (Thermo Fisher Scientific, Inc.). The RNAiso plus kit, Mini Best agarose gel DNA Purification kit, In-Fusion™ Advantage PCR Cloning kit, 3’-Full RACE Core Set Ver 2.0, 5’-Full RACE kit, DNA Ligation kit, Mini Best Plasmid Purification kit, molecular enzymes and primers used in this study were purchased from Takara Bio, Inc. 6 x-His antibody and horseradish peroxidase-Rabbit Antimouse IgG (H + L) were purchased from Proteintech Group, Inc. (Chicago, IL, USA). The bicinchoninic acid kit was from Nanjing KeyGen Biotech Co. (Nanjing, China). Ampicillin, chloramphenicol, tetracycline, kanamycin sulfate, isopropyl β-D-1-thiogalactopyranoside (IPTG) and phenylmethane sulfonyl fluoride were purchased from Tiangen Biotech Co., Ltd. (Beijing, China). The sypro orange protein gel stain was purchased from Sigma Aldrich (Merck KGaA, Darmstadt, Germany). The integrated potato culture medium, Luria Agar and Luria Broth were prepared in-house.

### 2.2 Cell line culture conditions

All the human cell lines used in this study were obtained from the Shanghai Cell Bank, Chinese Academy of Sciences (Shanghai, China). A panel of seven cell lines, including HeLa cervical carcinoma cells, HepG2 hepatoblastoma cells [reported mistakenly as hepatocellular carcinoma (Lopez-Terrada et al., 2009)], HL-60 adult acute myeloid leukemia cells, HCT-15 colorectal adenocarcinoma cells, SGC-7901 human gastric adenocarcinoma cells, SKOV-3 ovarian carcinoma cells and HaCaT human keratinocytes. HeLa, SKOV-3, HaCaT, HepG2 and SGC-7901 were grown in Dulbecco’s Modified Eagle’s Medium (Hyclone; GE Healthcare Life Sciences, Logan UT USA), while HCT-15 and HL-60 cells were grown in RPMI medium (Hyclone; GE Healthcare Life Sciences) containing 10% fetal bovine serum (TianJin Haoyang Biological Products Technology Co., Ltd., Tianjin, China) at 37°C in a humidified atmosphere containing 5% CO_2_.

### 2.3 Sequence retrieval and bioinformatics analysis of Lp16-PSP

The nucleotide and amino acid sequence of Lp16-PSP from *L. edodes* strain C_91-3_ were retrieved from the NCBI database (https://www.ncbi.nlm.nih.gov) (Accession nos. KF682441 and AHB81541, respectively). The *(Pfam)* database (https://pfam.xfam.org) was used for domain analysis of Lp16-PSP (Finn et al., 2016). The sequences of homologues of Lp16-PSP, i.e. endoribonuclease L-PSP from *Clostridium thermocellum*, Rut family protein from *Pyrococcus horikoshii,* protein mmf1 from *Saccharomyces cerevisiae,* putative endoribonuclease L-PSP from *Entamoeba histolytica,* ribonuclease UK114 from goat and ribonuclease from human were obtained in UniProt (http://www.uniprot.org) (UniProt nos. A3DJ68, O58584, P40185, C4LX79, P80601 and P52758, respectively). The alignment was accomplished by ClustalOmega (https://www.ebi.ac.uk/Tools/msa/clustalo/) (Sievers et al., 2011) and colored with ESPript 3.0 (http://espript.ibcp.fr/ESPript/ESPript/index.php) (Robert and Gouet, 2014). The secondary and three-dimensional (3D) structures of Lp16-PSP were predicted using JPred 4 (The Barton Group, School of Life sciences, University of Dundee, UK/ http://www.compbio.dundee.ac.uk/jpred4/index.html) (Drozdetskiy et al., 2015) and PHYRE2 Protein Fold Recognition Server (http://www.sbg.bio.ic.ac.uk/phyre2/html/page.cgi) (Kelley et al., 2015), respectively. The quality and reliability of the predicted models of Lp16-PSP were evaluated by Z-score, Root-Mean-Square Deviation (RMSD) value and Ramachandran plot analysis using the ProSA web server (https://prosa.services.came.sbg.ac.at/prosa.php) (Sippl, 1993, Wiederstein and Sippl, 2007), Dali server (http://ekhidna.biocenter.helsinki.fi/dali_server/start) (Holm and Rosenstrom, 2010) and RAMPAGE server (http://mordred.bioc.cam.ac.uk/~rapper/rampage.php) (Lovell et al., 2003), respectively. ExPASy server (https://web.expasy.org/compute_pi/) was used to establish the theoretical *pI* and molecular weight of Lp16-PSP (Bjellqvist et al., 1993, Bjellqvist et al., 1994).

### 2.4 cDNA synthesis, cloning and Lp16-PSP plasmid construction

The extraction of total RNA from *L. edodes* C_91-3_ mycelium and the synthesis of full-length cDNA was performed as previously described (Zhong et al., 2013). Oligo Primer Analysis Software v.6 (Molecular Biology Insights, Inc., Colorado Springs, CO, USA) was used to design the 3’ RACE and 5’ RACE primers based on the transcriptome sequence. The results of 3’ Full RACE and 5’ Full RACE were then sequenced and stitched and primers were designed accordingly. cDNA was synthesized by using Takara 3’-Full RACE Core Ver. 2.0. The cDNA was amplified by polymerase chain reaction (PCR) using the following primer sequences: Upstream primer *EcoR I,* 5’-GC<u>GAATTC</u>ACCAACAATGCATCCGGTG-3’ and downstream primer *Xho I 5*’-G<u>CTCGAG</u>ATTACAAGAGCGCTCAGTA-3’. The PCR reaction system (50 μl) consisted of the following: cDNA template, upstream primer, downstream primer, dNTP mixture (each 2.5 mM) 4 μl, 5 x PrimeSTAR buffer (Mg^2+^ plus) 10 μl, PrimeSTAR HS DNA Polymerase (2.5 U/μl) 0.5 μl (Takara Bio, Inc.) and nuclease-free water 33.5 μl. The reaction conditions were as follows: 30 cycles of 98°C for 10 sec, 56°C for 10 sec and 72°C for 1 min, followed by 72°C for 5 min and 4°C for 1 min. The ABI PRIMTM 3730XL DNA Sequencer (Thermo Fisher Scientific, Inc.) was used to confirm the amplified sequence. Amplified cDNA was run on 1 % agarose gel, purified by using MiniBEST Agarose Gel DNA Extraction Ver. 3.0 kit (Takara Bio, Inc.) and cloned into the *EcoRI/XhoI* sites of the pET32a (+) vector using the Infusion HD Cloning kit as per the manufacturer’s protocol (Takara Bio, Inc.). The in-fusion product was transformed into JM109 *E. coli* competent cells (Evans) and positive white colonies were confirmed by DNA sequencing. The plasmid pET32a *(+)-Lp16-PSP* was isolated and Rosetta-gami (DE3) *E. coli* was used as the expression strain.

### 2.5 Optimization of Lp16-PSP expression conditions and recovery of inclusion bodies

For the preliminary expression of Lp16-PSP, Luria Broth (10 g/l Tryptone, 5 g/l Yeast extract and10 g/1 NaCl, pH 7.0) containing appropriate antibiotics (chloramphenicol 34 μg/ml, tetracycline 12.5 μg/ml, kanamycin sulphate 15 μg/ml and ampicillin 100 μg/ml) was used to grow a single colony of transformants at 37°C overnight in orbital shaker. A total of 1 ml of the overnight culture was refreshed in 10 ml of LB medium and grown at 37°C in an orbital shaker until the optical density at 600 nm (OD_600_) reached 0.6 as measured on a microplate reader. To induce fusion protein expression, IPTG was added at a final concentration of 0.05 mM and cells were incubated for 3 h at 37°C on an orbital shaker. An uninduced control culture was also set up. Bacteria were subsequently centrifuged at 3,824 x g at 4°C for 5 min. The bacterial cell pellet was washed once with PBS pH 7.4 and resuspended in lysis buffer (Table 1). The mixture was shaken at 4°C for 1 h, sonicated for 10 cycles of 20 W for 1 min with 1 min breaks on ice. The lysed bacterial suspension was centrifuged at 15,297 x g for 20 min at 4°C. The supernatant and pellet were collected and samples underwent western blotting for His-Tagged protein. Briefly, equal volume of samples (10 μ1) were separated by 12% SDS-PAGE and transferred to polyvinylidene difluoride membranes, followed by blocking with 5% skimmed milk at room temperature for 1 h. After blocking, membranes were incubated with anti-6 x His antibody (66005-1-Ig; 1:1,000; Proteintech, Inc.) at 4°C overnight. Membranes were washed with TBS-T three times and incubated with second antibody goat anti-mouse IgG (SA00001-1; 1:6000, Proteintech, Inc.) at room temperature for 1 h. After washing three times with TBS-T, blots were developed using an ECL Ultra kit (New Cell & Molecular Biotech Co. Ltd, China) and images were captured using a ChemiDocTM XRS + Imager (Bio-Rad Laboratories, Inc., Hercules, CA, USA).

**Table 1.**
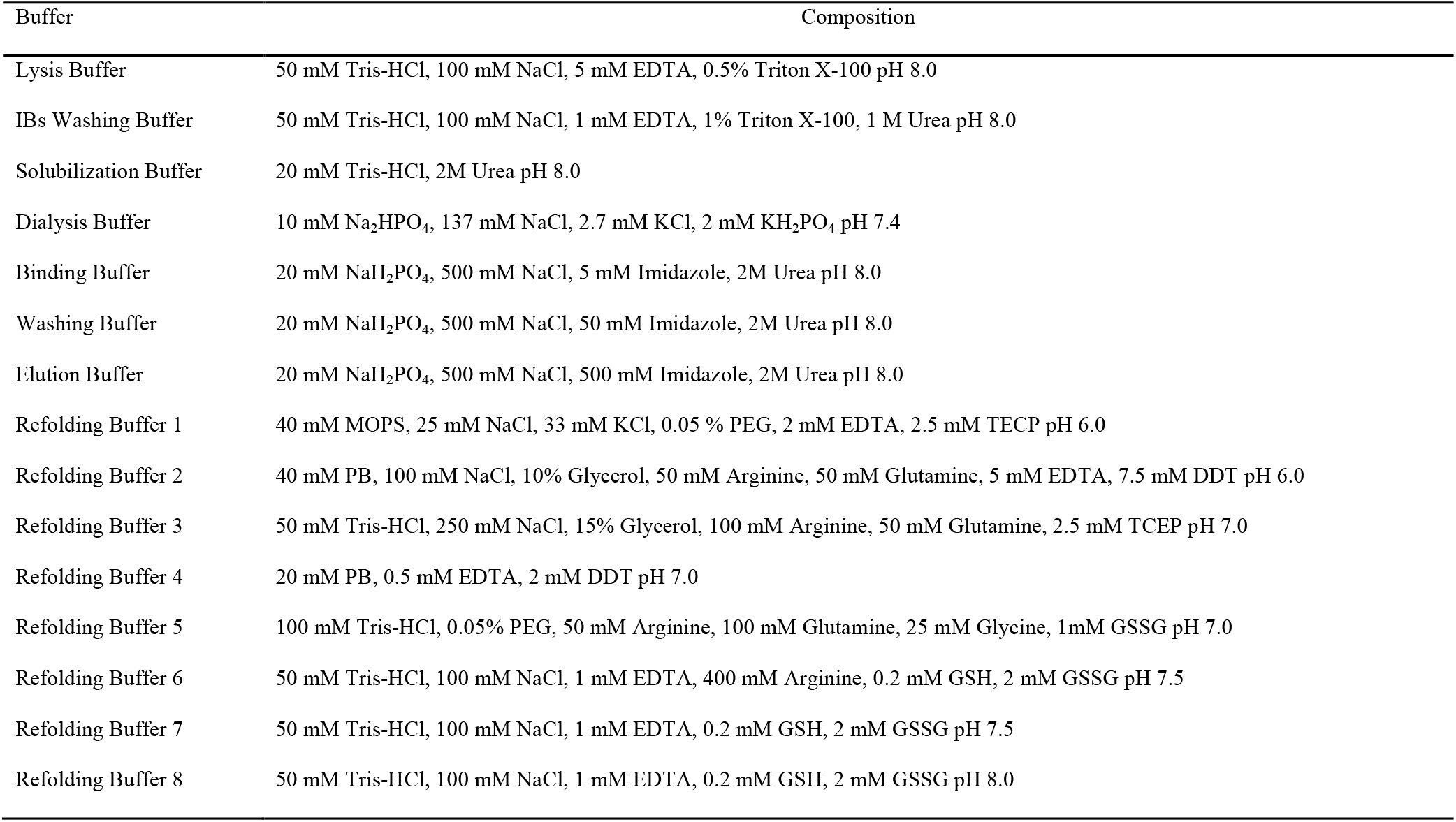
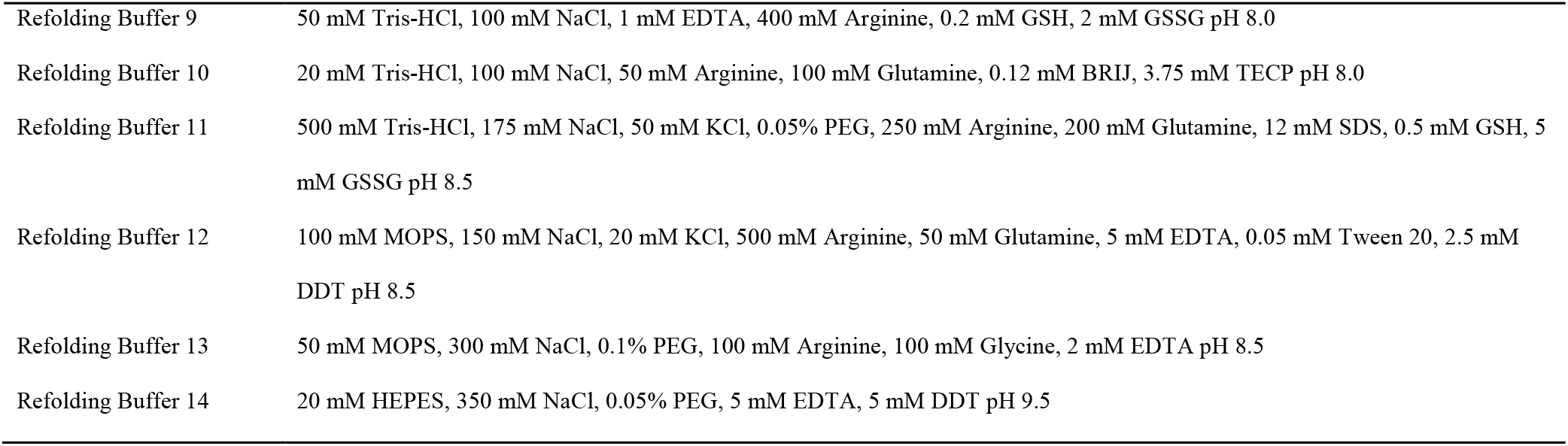
Buffers used in the present study

| Buffer | Composition |
| --- | --- |
| Lysis Buffer | 50 mM Tris-HCl, 100 mM NaCl, 5 mM EDTA, 0.5% Triton X-100 pH 8.0 |
| IBs Washing Buffer | 50 mM Tris-HCl, 100 mM NaCl, 1 mM EDTA, 1% Triton X-100, 1 M Urea pH 8.0 |
| Solubilization Buffer | 20 mM Tris-HCl, 2M Urea pH 8.0 |
| Dialysis Buffer | 10 mM Na<sub>2</sub>HPO<sub>4</sub>, 137 mM NaCl, 2.7 mM KCl, 2 mM KH<sub>2</sub>PO<sub>4</sub> pH 7.4 |
| Binding Buffer | 20 mM NaH<sub>2</sub>PO<sub>4</sub>, 500 mM NaCl, 5 mM Imidazole, 2M Urea pH 8.0 |
| Washing Buffer | 20 mM NaH<sub>2</sub>PO<sub>4</sub>, 500 mM NaCl, 50 mM Imidazole, 2M Urea pH 8.0 |
| Elution Buffer | 20 mM NaH<sub>2</sub>PO<sub>4</sub>, 500 mM NaCl, 500 mM Imidazole, 2M Urea pH 8.0 |
| Refolding Buffer 1 | 40 mM MOPS, 25 mM NaCl, 33 mM KCl, 0.05 % PEG, 2 mM EDTA, 2.5 mM TECP pH 6.0 |
| Refolding Buffer 2 | 40 mM PB, 100 mM NaCl, 10% Glycerol, 50 mM Arginine, 50 mM Glutamine, 5 mM EDTA, 7.5 mM DDT pH 6.0 |
| Refolding Buffer 3 | 50 mM Tris-HCl, 250 mM NaCl, 15% Glycerol, 100 mM Arginine, 50 mM Glutamine, 2.5 mM TCEP pH 7.0 |
| Refolding Buffer 4 | 20 mM PB, 0.5 mM EDTA, 2 mM DDT pH 7.0 |
| Refolding Buffer 5 | 100 mM Tris-HCl, 0.05% PEG, 50 mM Arginine, 100 mM Glutamine, 25 mM Glycine, 1mM GSSG pH 7.0 |
| Refolding Buffer 6 | 50 mM Tris-HCl, 100 mM NaCl, 1 mM EDTA, 400 mM Arginine, 0.2 mM GSH, 2 mM GSSG pH 7.5 |
| Refolding Buffer 7 | 50 mM Tris-HCl, 100 mM NaCl, 1 mM EDTA, 0.2 mM GSH, 2 mM GSSG pH 7.5 |
| Refolding Buffer 8 | 50 mM Tris-HCl, 100 mM NaCl, 1 mM EDTA, 0.2 mM GSH, 2 mM GSSG pH 8.0 |
| Refolding Buffer 9 | 50 mM Tris-HCl, 100 mM NaCl, 1 mM EDTA, 400 mM Arginine, 0.2 mM GSH, 2 mM GSSG pH 8.0 |
| Refolding Buffer 10 | 20 mM Tris-HCl, 100 mM NaCl, 50 mM Arginine, 100 mM Glutamine, 0.12 mM BRIJ, 3.75 mM TECP pH 8.0 |
| Refolding Buffer 11 | 500 mM Tris-HCl, 175 mM NaCl, 50 mM KCl, 0.05% PEG, 250 mM Arginine, 200 mM Glutamine, 12 mM SDS, 0.5 mM GSH, 5 mM GSSG pH 8.5 |
| Refolding Buffer 12 | 100 mM MOPS, 150 mM NaCl, 20 mM KCl, 500 mM Arginine, 50 mM Glutamine, 5 mM EDTA, 0.05 mM Tween 20, 2.5 mM DDT pH 8.5 |
| Refolding Buffer 13 | 50 mM MOPS, 300 mM NaCl, 0.1% PEG, 100 mM Arginine, 100 mM Glycine, 2 mM EDTA pH 8.5 |
| Refolding Buffer 14 | 20 mM HEPES, 350 mM NaCl, 0.05% PEG, 5 mM EDTA, 5 mM DDT pH 9.5 |

In order to optimize the Lp16-PSP expression conditions, three independent parameters were validated: Incubation temperature, concentration of inducer and post-induction incubation time. Rosetta-gami (DE3) *E. coli* bearing pET32a (+) - *Lp16-PSP* was grown in LB medium overnight at 37°C. The overnight culture was diluted in fresh LB media supplemented with chloramphenicol (34 μg/ml), tetracycline (12.5 μg/ml), kanamycin sulphate (15 μg/ml) and ampicillin (100 μg/ml) at a ratio of 1:10 and cultured at 37°C till the OD_600_ reached 0.6. The culture was subsequently induced with IPTG (final concentration 1 mM) and incubated for 3 h at 20, 30 or 37°C. To optimize the inducer concentration, midexponential-phase cultures were induced with different concentrations of IPTG (0.05-1 mM) and incubated for 3 h at 37°C. To optimize the post-induction incubation period, mid-exponential phase culture was induced with 0.5 mM IPTG and incubated at 37°C for 1-6 h. Bacteria were harvested and centrifuged at 3,824 x g at 4°C for 5 min and lysis was performed as above. The lysed bacterial suspension centrifuged at 15,294 x g for 20 min at 4°C. The supernatant and pellet were collected and proteins (10 μl) were separated by 12% SDS-PAGE analysis. Protein bands were quantified using Image Lab Software v.4.0.1 (Bio-Rad Laboratories, Inc.).

### 2.6 Solubilization of inclusion bodies in different solvents with different pH and temperatures

Following lysis, protein pellets were washed three times with washing buffer (Table 1). In order to remove contaminating detergents, pellets were washed with PBS and centrifuged at 15,294 x g for 20 min at 4°C. The washed inclusion bodies were subsequently resuspended in Milli Q water. The homogeneous suspension of Lp16-PSP inclusion bodies (~10 mg/ml) was solubilized in various solubilization buffers (PBS pH 8.0, 20 mM sodium phosphate pH 8.0, 20 mM potassium phosphate pH 8.0, 20 mM Tris-HCl pH 8.0 and deionized water pH 8.0) containing 2 M urea. Suspensions were subsequently frozen at – 20°C and thawed at room temperature. In order to observe the effect of pH on the solubility of Lp16-PSP inclusion bodies, a homogenous suspension of Lp16-PSP inclusion bodies (~10 mg/ml) was solubilized in 20 mM Tris-HCl ranging from pH 5-9 with 2 M urea. Suspensions were frozen at – 20°C and then thawed at room temperature. Homogeneous suspensions of Lp16-PSP inclusion bodies (~10 mg/ml) were solubilized in 20 mM Tris-HCl pH 8.0 at different temperatures 20, 30, −20, −40 or −80°C, or in liquid nitrogen (−196°C). All samples after were centrifuged at 15,294 x for 20 min at 4°C and supernatants were collected (Qi et al., 2015). A total of 10 μl of the collected supernatants were analyzed by SDS-PAGE and protein bands were quantified using Image Lab Software v.4.0.1 (Bio-Rad Laboratories, Inc.).

### 2.6 Purification of recombinant Lp16-PSP protein

The recombinant Lp16-PSP protein has a tag comprising of 6 histidines, and so purification was performed using Ni-magnetic beads (Tomos Biotools Shanghai Co., Ltd., Shanghai, China) according to the manufacturer’s protocol. Briefly, beads were washed three times with deionized water and binding buffer (Table 1). The solubilized protein supernatant was subsequently agitated with washed Ni-beads for 1 h at 4°C. Unbound proteins were then removed using wash buffer (Table 1) three times at 4°C. Following washing, samples were eluted with elution buffer (Table 1). The concentration of the purified protein was determined using the Bradford method (Kruger, 1994) and proteins (10 μl) were separated by 12% SDS-PAGE subjected to western blotting as described above.

### 2.7 Differential scanning fluorometry (DSF) guided refolding of Lp16-PSP

Purified inclusion bodies were refolded in 1.5 ml Eppendorf tubes by diluting the protein in a variety of refolding buffers in a ratio of 1:20 (Table 1). Tubes were stored at 4°C overnight and precipitates were removed by centrifugation at 3,700 x g for 20 min at 4°C. DSF assays were performed (Biter et al., 2016) using the StepOne^TM^ RealTime PCR machine (48 wells) with the following thermal profile: Step 1, 100% ramp rate, 25°C for 2 min; Step 2, 1% ramp rate, 99°C for 2 min. Together with refolded samples, denatured Lp16-PSP and no protein control samples were also run. The quantitative PCR machine-generated data file was then analyzed by using protein thermal shift software v1.3 (Thermo Fisher Scientific, Inc.). Boltzmann Tm values and Boltzmann fit were then taken into consideration. The optimized refolding buffer was then used for routine refolding of Lp16-PSP protein using the direct dilution method (Vallejo and Rinas, 2004, Singh and Panda, 2005, Lilie et al., 1998).

### 2.8 Dialysis and concentration of Lp16-PSP protein

The refolded protein was placed in a dialysis membrane (MWCO 14 kDa, Biosharp Hefei, China) and was subjected to dialysis with ten volumes of dialysis buffer at 4°C for 12 h with two buffer changes (Table 1). The dialyzed protein was concentrated using PEG 20,000. The final protein product was quantified using a bicinchoninic acid assay and assessed using 12% SDS-PAGE.

### 2.9 Cell viability assay

A cell viability assay was performed using Cell Counting Kit-8 (CCK-8); Biotool) following the manufacturer’s protocol. Briefly, 5×10^3^ cells of adherent cell lines and 1×10^4^ cells of the non-adherent cell lines were seeded in 96 well plates and grown overnight at 37°C. Cells were subsequently treated with 0, 12.5, 25, 50, 100, 150 or 200 μg/ml Lp16-PSP and incubated at 37°C for 24 or 48 h. A volume of 10 μl CCK-8 reagent was added to each well and cells were incubated at 37°C for 4 h. The absorbance was measured on an ELISA plate reader at 450 nm. The cell viability was calculated as follows (He et al., 2015): Cell viability (%) = A_Experimental group_/A_Control group_ x 100. Where, “A” is the absorbance at 450 nm.

### 2.10 Phase contrast imaging

HL-60 cells were seeded in 6-well plates and incubated overnight at 37°C. Cells were subsequently washed with PBS once and grown for 48 h in RPMI medium with and without Lp16-PSP (0, 50, 100 and 150 μg/ml). Phase contrast images were captured using fluorescence microscopy at 40 x magnification.

### 2.11 Statistical analysis

GraphPad Prism 5 software (GraphPad, Inc., La Jolla, CA, USA) was used for statistical analysis. One-way analysis of variance was used followed by Tukey’s comparison test. P<0.05 was considered to indicate a statistically significant difference. The half maximal inhibitory concentration (IC_50_) of Lp16-PSP was calculated using linear regression on Microsoft Excel 2007 (Microsoft Corporation, Redmond, WA, USA).

## 3.0 Results

### 3.1 Bioinformatics analysis of Lp16-PSP protein

The nucleotide and amino acid sequences of Lp16-PSP from *L. edodes* strain C_91-3_ were retrieved using the NCBI database with the accession numbers KF682441 and AHB81541, respectively and analyzed using several bioinformatics tools. The specific ‘Domain’ protein region gives the protein a unique function or interaction, contributing to defining its overall role (Zhang et al., 2012). In the present study, the *Pfam* database was used for conserved domain analysis of Lp16-PSP and it was revealed that the Lp16-PSP protein contains an endoribonuclease L-PSP domain (**Fig. 1A**) and belongs to the YjgF/YER057c/UK114 protein superfamily. The multiple sequence alignment of Lp16-PSP with its homologs revealed 12 invariantly conserved residues of the YjgF/YER057c/UK114 family (**Fig. 1B**). The overall structure of Lp16-PSP has an α+β fold, arranged as βββαβαββ, which is typically referred to as chorismate mutase-like fold (Volz, 1999, Sinha et al., 1999a). The order of the strands in the sheet is β_1_β_2_β_3_β_6_β_4_β_5_ and all are antiparallel, aside from β_4_β_5_, which are parallel to each other. The 3D structure of Lp16-PSP was predicted using the PHYRE 2 server (**Fig. 2A**). The Z-score value of Lp16-PSP was >2, while the RMSD value was <1. As per the Ramachandran plot, 95.9% of Lp16-PSP residues were in the favored region, which indicates that the 3D models of Lp16-PSP were correct (**Fig. 2B**). The theoretical *pI* and molecular weight of Lp16-PSP, as determined by Expasy’s server, were 9.05 and 14.3 kDa, respectively.

**Figure 1.**
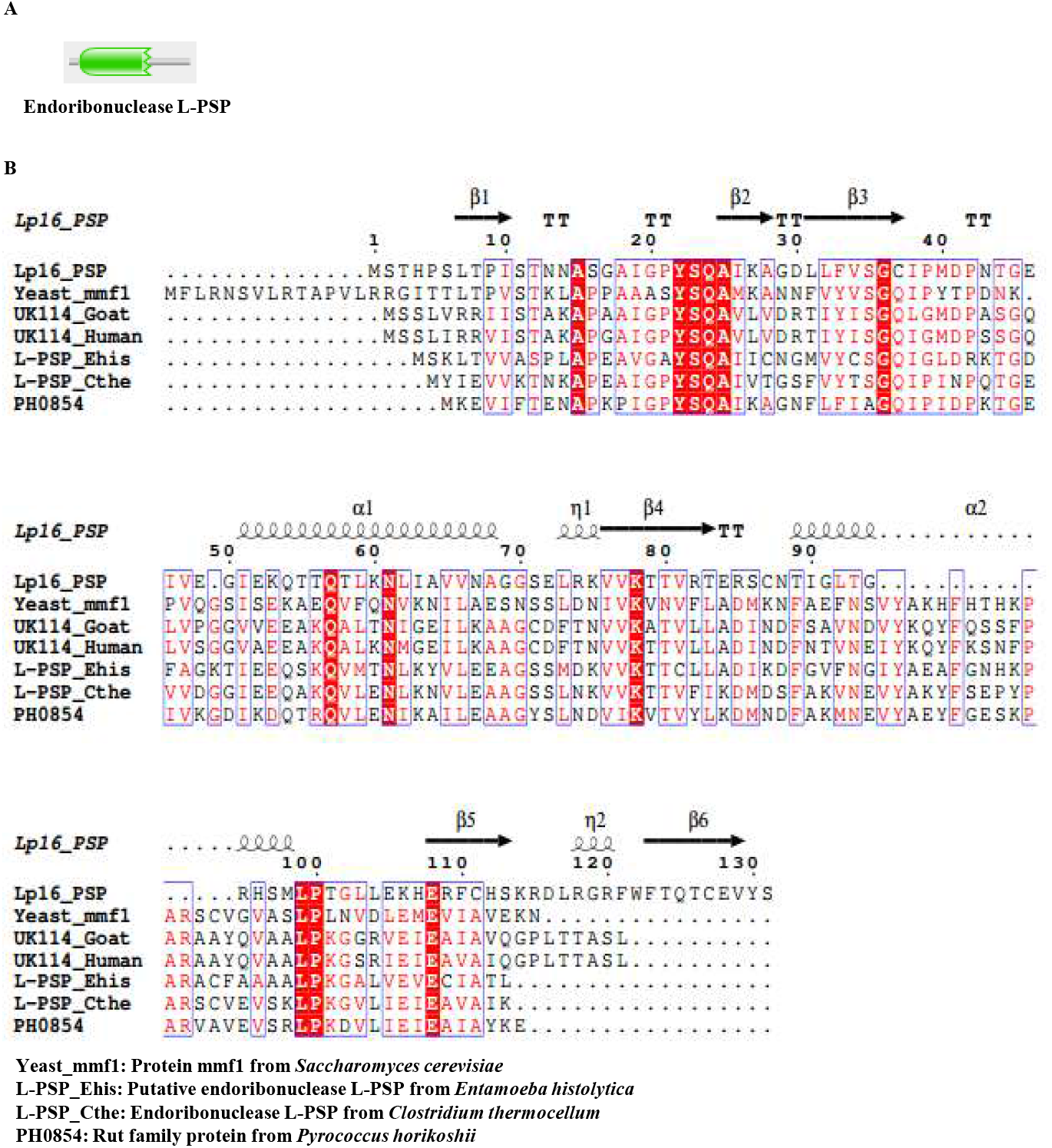
Domain analysis and multiple sequence alignment of Lp16-PSP. **(A)** Domain analysis of Lp16-PSP, revealing the presence of an endoribonuclease L-PSP domain. **(B)** Multiple sequence alignment of Lp16-PSP with other members of the YjgF/ YER057c/UK114 protein family. Helices, and arrows represent α-helix and β-sheets, respectively. The 12 conserved residues are highlighted in red color. Lp16-PSP, Latcripin-16.

### 3.2 Cloning, transformation and expression of Lp16-PSP in Rosetta-gami (DE3) E. coli

Lp16-PSP cDNA was cloned from the *L. edodes* strain C_91-3_ using 3’ RACE and 5’ RACE techniques. The DNA sequence encoding Lp16-PSP was cloned downstream of a T7 promoter in the expression vector pET32a (+) (**Fig. 2C**). The sequence was verified by DNA sequencing and the resulting plasmid was transformed into Rosetta-gami (DE3) *E. coli* for protein expression. Prominent protein bands were observed in the supernatant and pellet of the induced samples following lysis of *E. coli* cells harboring pET32a *(+)-Lp16-PSP* plasmid, with a molecular mass of 32 kDa. The results also demonstrated that a major portion of the Lp16-PSP was expressed as an inclusion body.

**Figure 2.**
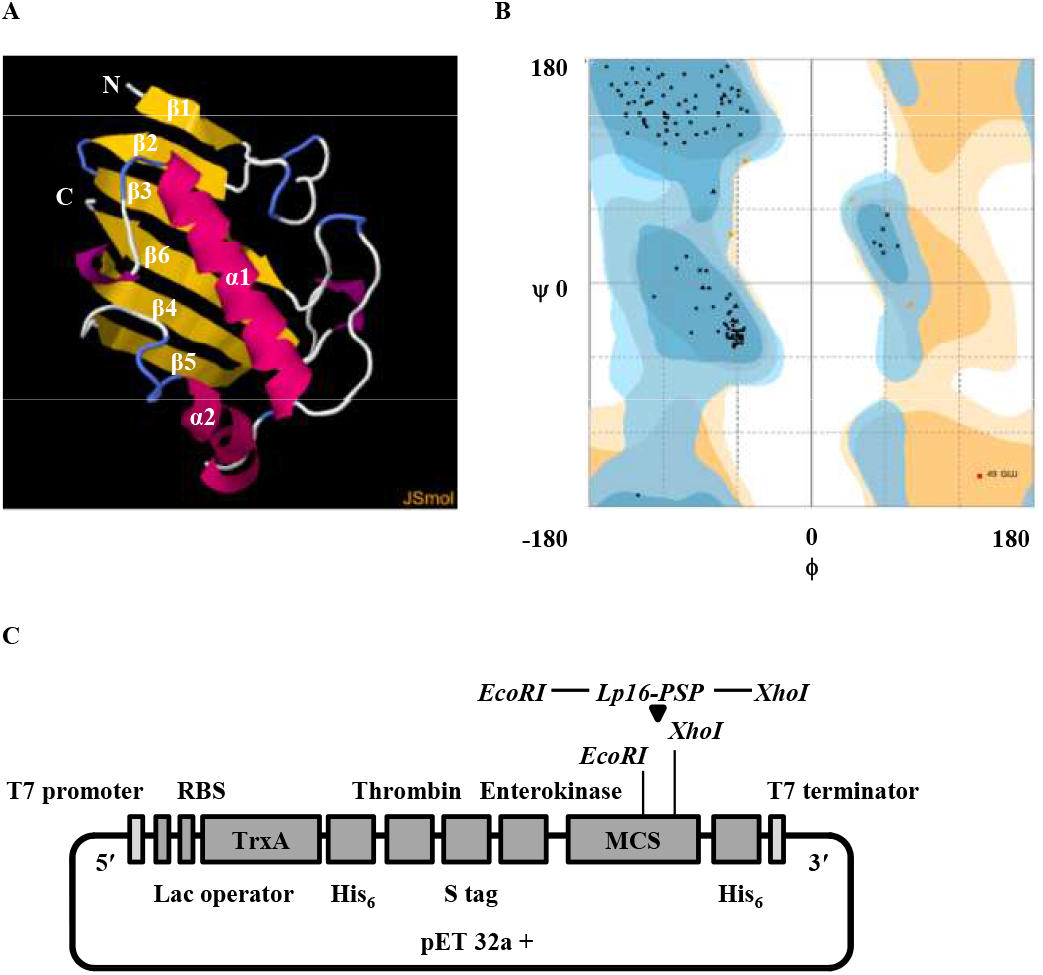
Three-dimensional structure and cloning of Lp16-PSP. **(A)** Three-dimensional structure of Lp16-PSP, which is homologous to yeast mmf1 protein (PDB I.D c3quwA) with 100% confidence and 93% coverage. All β-sheets are anti-parallel, aside from β4 and β5, which are parallel to each other. **(B)** Ramachandran plot demonstrating the quality and reliability of the Lp16-PSP three-dimensional structure. The number of residues in favored, allowed and outlier regions are 95.9, 3.3 and 0.8%, respectively. **(C)** Schematic representation of pET32a (+) plasmid with Lp16-PSP inserted at *EcoRI* and *XhoI* sites. Lp16-PSP, Latcripin-16; RBS, ribosome binding site; Lac, lactose; His6, six histidine tag; MCS, multiple cloning site; TrxA, thioredoxin A; S tag, solubility tag.

Following the successful initial expression and confirmation of Lp16-PSP, the optimal expression parameters were investigated. Various temperatures, IPTG concentrations and induction times were assessed. The maximal expression of Lp16-PSP was observed at 37°C and nearly all of the protein was located in the pellet as an inclusion body (**Fig. 3A**). *E. coli* cells grow over a wide range of temperatures (15-42°C), however the growth rate increased proportionally with temperature increases between 20 and 37°C. It has previously been reported that, at 23-42°C, the number of ribosomes per cell and level of rRNA remains constant (Malik et al., 2016). In fact, the peptide chain elongation rate increases with increased temperatures, resulting in increased protein synthesis (Farewell and Neidhardt, 1998). Therefore, Lp16-PSP synthesis was faster at higher temperatures due to an increase in plasmid copy number, transcription rate and peptide chain elongation rate. The next step was to optimize the concentration of IPTG. No significant differences in Lp16-PSP expression were observed between different IPTG concentrations (**Fig. 3B**). However, treatment with 0.5 or 1 mM of IPTG resulted in a slightly higher expression of Lp16-PSP compared with lower concentrations. As such, 0.5 mM IPTG was used for subsequent experiments. To assess the induction time, kinetic studies were performed at 37°C, samples were taken every hour and analyzed using 12% SDS-PAGE. The maximal expression of Lp16-PSP was observed at 4 h; thereafter, the expression remained constant until 6 h (**Fig. 3C**). Based on these results, the optimal expression conditions of Lp16-PSP were selected as 37°C with 0.5 mM IPTG for 4 h (**Fig. 3D**). Cells were subsequently collected, lysed, centrifuged and washed to obtain the pelleted crude inclusion bodies, which contained Lp16-PSP and were then used for the solubilization and refolding purposes.

**Figure 3.**
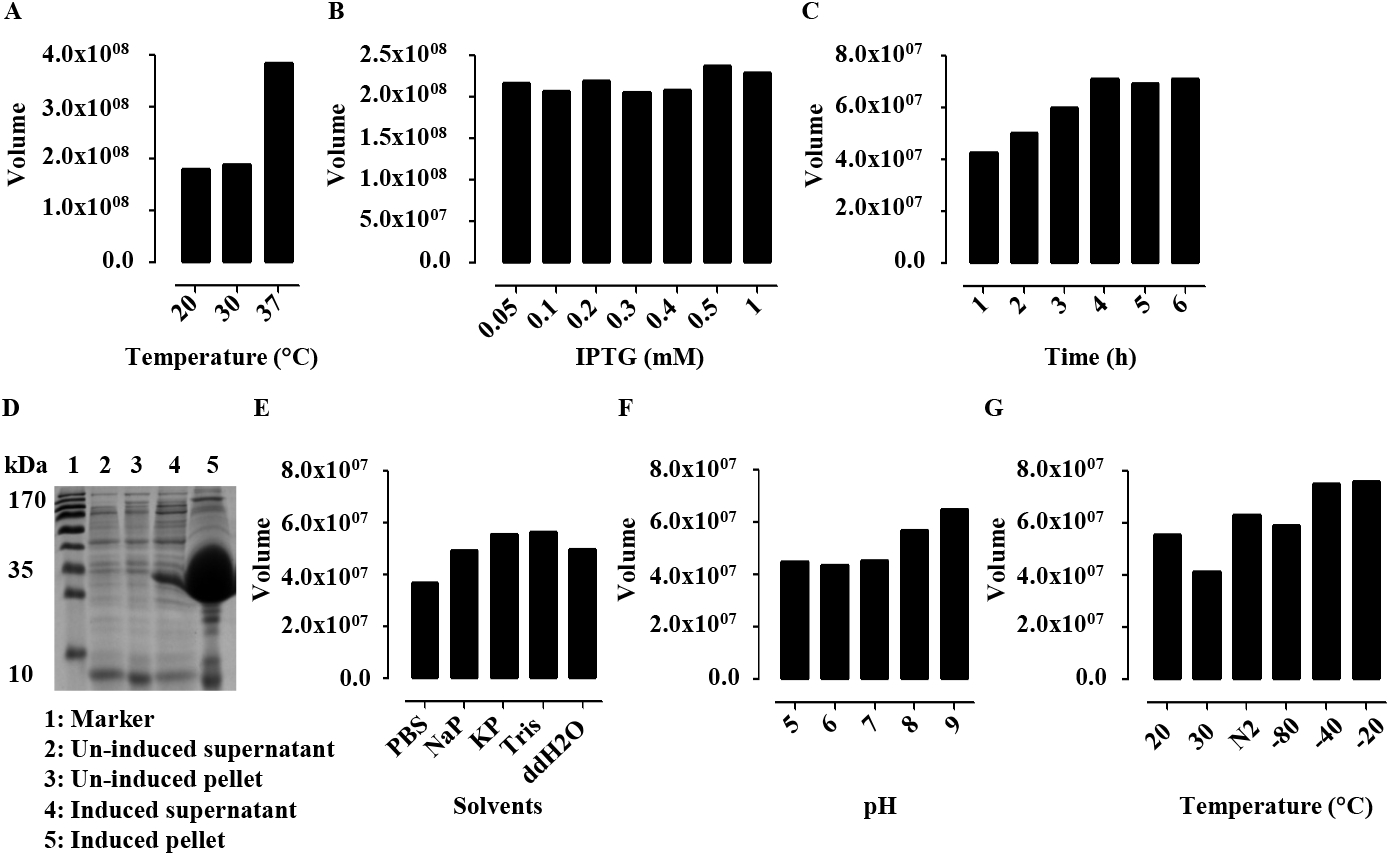
Efficient production of Lp16-PSP. Effects of **(A)** temperature, **(B)** IPTG concentration and **(C)** induction time on Lp16-PSP production. **(D)** SDS-PAGE analysis of Lp16-PSP expression under optimized conditions. Effects of **(E)** different solvents, **(F)** pH and **(G)** temperature on the solubility of Lp16-PSP inclusion bodies. Lp16-PSP, Latcripin-16; IPTG, isopropyl β-D-1-thiogalactopyranoside.

### 3.3 Solubilization of Lp16-PSP IBs

In order to achieve the maximum amount of Lp16-PSP in solution form and to prevent the introduction of deleterious modifications, the solubility of Lp16-PSP was studied using different solvents. The results indicate that various buffers containing 2 M urea are able to efficiently solubilize Lp16-PSP protein from inclusion bodies; however, the maximum solubility was observed with 20 mM Tris-HCl (pH 8.0); **Fig. 3E**). Furthermore, Lp16-PSP inclusion bodies were solubilized in 20 mM Tris-HCl buffer at different pHs 5-9 and quantitatively monitored (**Fig. 3F**). It was observed that increased pH resulted in increased solubilization of Lp16-PSP. The effect of solubilization temperature was assessed and maximum solubility was observed at −20 and −40°C. However, to prevent the protein from harsh freeze-thaw shock, − 20 °C was selected for further experiments (**Fig. 3G**).

### 3.4 Purification and refolding of a recombinant Lp16-PSP protein

Under denaturing conditions, Lp16-PSP was purified using Ni magnetic beads. Binding of the Lp16-PSP with Ni magnetic beads was performed at 4°C with constant shaking. In order to prevent non-specific binding of the histidine-rich proteins, 500 mM NaCl and 5 mM imidazole were added in the binding buffer. Beads were washed three times with washing buffer containing 50 mM imidazole to elute the non-specifically bound proteins. Finally, Lp16-PSP was eluted using 500 mM imidazole in the elution buffer. At this stage, protein was almost pure; the bands of Lp16-PSP appeared at 32 kDa and eluted His-tagged Lp16-PSP was further confirmed by western blotting (**Fig. 4A**).

**Figure 4.**
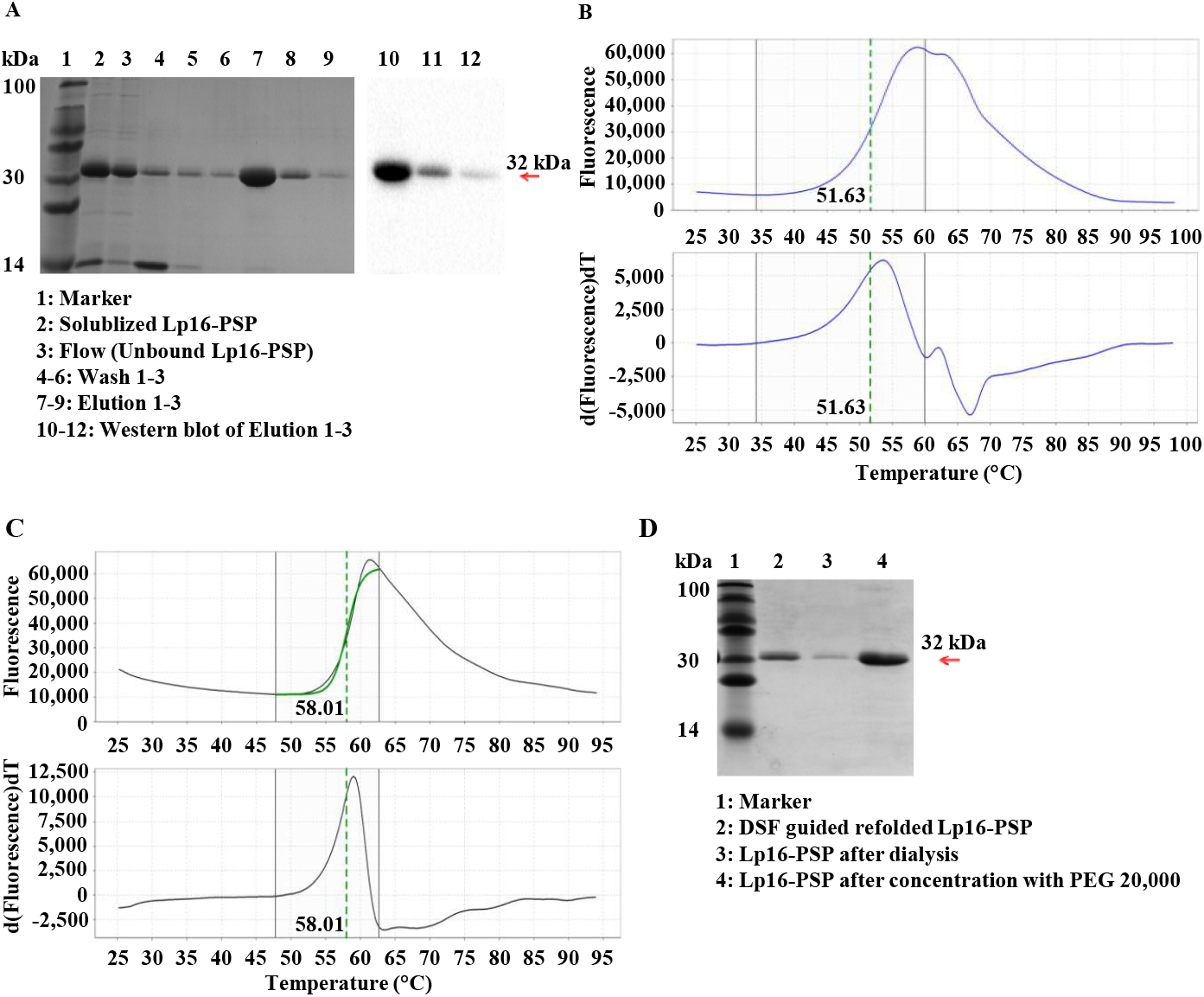
Purification and DSF guided refolding of Lp16-PSP. (A) SDS-PAGE and western blotting of Lp16-PSP following purification using Ni magnetic beads. **(B)** DSF of denatured Lp16-PSP protein. Straight black lines represent the region of analysis and dotted green lines represent the Boltzmann Tm value (51.63). **(C)** DSF-guided refolding of Lp16-PSP protein straight black lines represent the region of analysis and dotted green lines represent the Boltzmann Tm value (58.01). **(D)** SDS-PAGE of the finalized Lp16-PSP protein. DSF, Differential Scanning Fluorometry; Lp16-PSP, Latcripin-16.

*E. coli* offers rapid and low-cost protein production; however, the majority of the proteins expressed are in the form of inclusion bodies (Graslund et al., 2008). Although proteins in these inclusion bodies are relatively pure (Burgess, 2009), they are misfolded, amorphous protein aggregates (Baneyx and Mujacic, 2004) that are biologically inactive. Therefore, proper *in vitro* refolding of the expressed protein is performed to obtain the bioactive form. In the present study, mild solubilization conditions were used to preserve the native-like structure of Lp16-PSP for further *in vitro* refolding.

DSF screening was used to determine and optimize the refolding conditions. Since the Anfnsen hypothesis states that “the native state of a folded protein is also the most thermodynamically stable state” (Anfinsen, 1973), it was speculated that Lp16-PSP would have the highest Tm value with the buffer responsible for proper refolding. Different buffers ranging in pH from 6-9.5 were used in the present study (**Table 1**) and the values of Tm x peak height were determined. It was demonstrated that denatured Lp16-PSP produced a thermal melting transition of Tm 51.63°C (**Fig. 4B**). In contrast, refolding buffer resulted in the highest Tm value of 58.01°C, indicating proper refolding Lp16-PSP (**Fig. 4C**). It is known that the mild solubilization process is advantageous as it prevents the complete unfolding of the protein’s native-like structure (Khan et al., 1998). Following refolding with validated refolding buffer, the refolded protein was extensively dialyzed and concentrated using PEG 20,000. This concentrated protein was qualitatively (**Fig. 4D**) and quantitatively examined and was used for subsequent biological assays.

### 3.4 Lp16-PSP showed selective cytotoxicity against human cancerous and non-cancerous cell lines

The cytotoxic activity of Lp16-PSP was evaluated using a CCK-8 assay. HL-60, HepG2, HeLa, SGC-7901, SKOV-3, HCT-15 and HaCaT cell lines were treated with several concentrations of 0, 12.5, 25, 50, 100, 150 or 200 μg/ml Lp16-PSP for 24 or 48 h and cell viability was determined. The IC_50_ of Lp16-PSP for each cell line is presented in **Fig. 5A**, while the cytotoxicity data of HL-60 cell line is presented in **Fig. 5B**. All cancer cell lines (HL-60, HepG2, HeLa, SGC-7901, SKOV-3 and HCT-15) were sensitive to Lp16-PSP, whereas the non-cancerous HaCaT cell line was less sensitive to Lp16-PSP.

**Figure 5.**
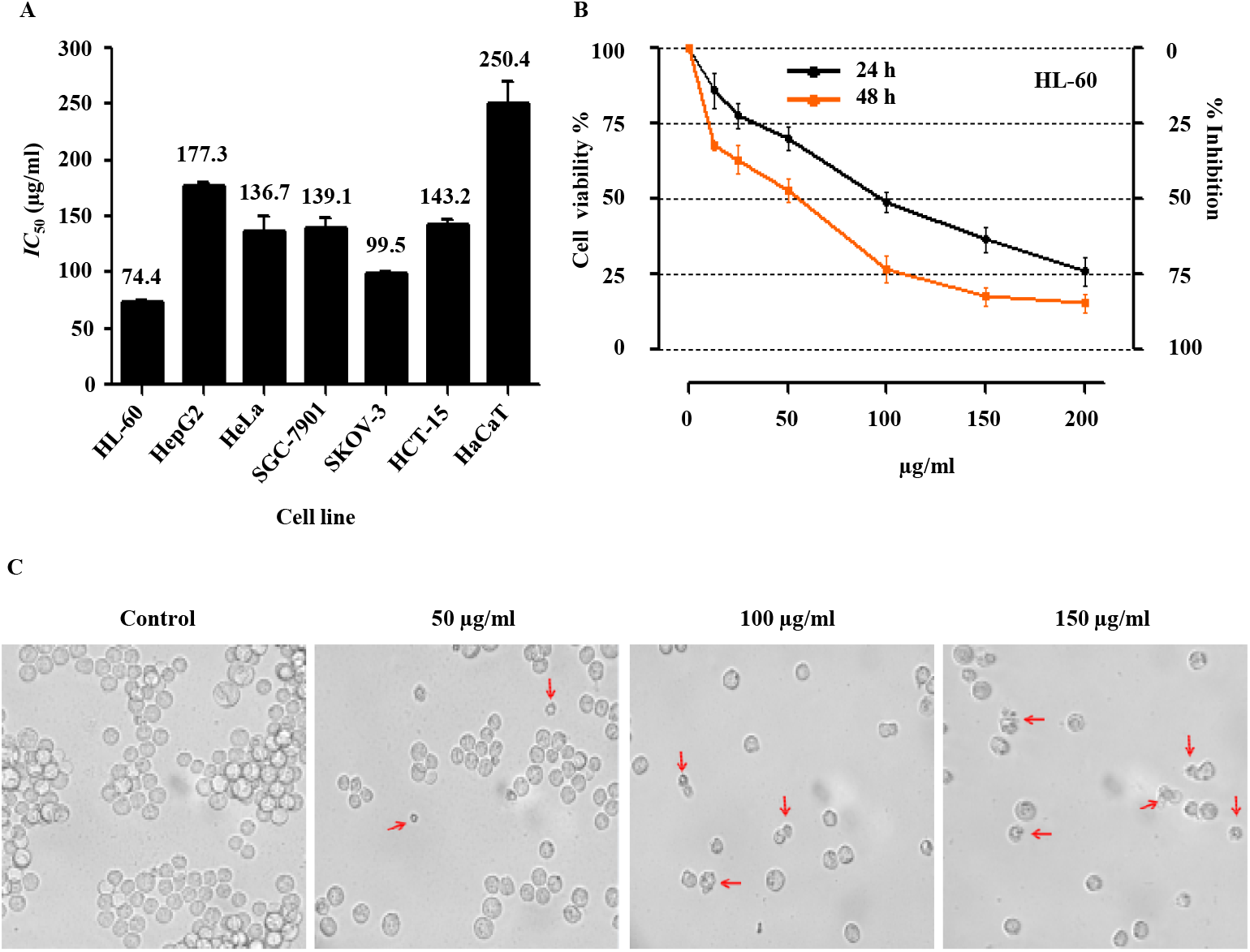
Biological activity of Lp16-PSP. **(A)** IC_50_ values of Lp16-PSP in human cancerous and non-cancerous cell lines following 48 h incubation. Data are presented as the mean ± standard error of the mean. **(B)** Viability of HL-60 cells treated with Lp16-PSP for 24 or 48 h as assessed using a Cell Counting Kit-8 assay. Data are presented as the mean ± standard deviation. **(C)** Phase contrast images of HL-60 cells following treatment with Lp16-PSP for 48 h. Red arrows indicate small cells and cellular bleeding. Magnification, x40. Lp16-PSP, Latcripin-16; IC_50_, half maximal inhibitory concentration.

Phase contrast images of HL-60 cells were captured following treatment (**Fig. 5C**). There was an obvious change in cell volume and size between Lp16-PSP treated HL-60 cells and control cells. The reduction in cell size, abnormal morphological changes and cell bursting suggest that cells treated with Lp16-PSP underwent apoptosis or are dead. These findings demonstrate that Lp16-PSP may act as a broadspectrum anticancer agent with selective cytotoxicity.

## 4.0 Discussion

The Lp16-PSP gene isolated from *L. edodes* C_91-3_ is an endoribonuclease L-PSP and a member of highly conserved YjgF/YER057c/UK114 superfamily. Members of the YjgF/YER057c/UK114 protein family have previously been reported to be endoribonucleases, translation inhibitors and antiviral agents (Morishita et al., 1999, Manjasetty et al., 2004, Su et al., 2015). To combat cancer, gene expression in cancer cells can be controlled at different levels. Agents that target RNA are preferred over agents targeting DNA due to the genotoxic effects of the DNA targeting agents (Gurova, 2009) that may lead to a new and resistant cancer. There has been progress research into antitumor RNases and a number of natural and recombinant RNases has been reported to have anticancer abilities (Arnold and Ulbrich-Hofmann, 2006, Mutti and Gaudino, 2008, Saxena et al., 2003, Lee and Raines, 2008, Castro et al., 2011). In the present study, it was speculated that Lp16-PSP, as a member of the YjgF/YER057c/UK114 family, may exert its anticancer activity by targeting and degrading RNA and inhibiting translation.

In order to evaluate the anticancer activity of Lp16-PSP, the Lp16-PSP gene was cloned into a prokaryotic expression vector pET 32a (+) and expressed as His-tagged fusion protein in *E. coli* Rosetta gami *(DE3)* under optimized conditions. Because the majority of the protein was expressed as inclusion bodies, the solubilization of Lp16-PSP was optimized using various solvents at different pH and temperatures. Pure Lp16-PSP protein was recovered via the routine affinity purification method. Solubilization with mild solubilization buffer containing 2 M urea at pH 8.0 by freeze-thaw method did not result in the complete denaturation of Lp16-PSP and thus refolding was achieved easily. Following the recovery of purified Lp16-PSP, its anticancer activity was investigated against a panel of human cancerous and non-cancerous cell lines. The IC_50_ value of Lp16-PSP was lower in cancerous cell lines compared with non-cancerous cell lines, highlighting the selective toxic effect of Lp16-PSP. In addition, adult acute myeloid leukemia (HL-60 cells) were the most sensitive cell lines assessed in the present study. Morphological changes in HL-60 cells following treatment with Lp16-PSP were indicative of apoptosis.

The results of the present study suggest that Lp16-PSP may serve as a potential anticancer agent; however, these results are preliminary and the effects of Lp16-PSP have only been investigated *in vitro.* Future studies should use animal models to assess the anticancer activity of Lp16-PSP *in vivo.* The cytotoxic Lp16-PSP doses used in the present study were very high, and so future studies should aim to identify strategies that may improve the delivery of Lp16-PSP in *in vitro* and *in vivo* models (Walev et al., 2001, Zochowska et al., 2009, Kaczmarczyk et al., 2011, Erazo-Oliveras et al., 2014, Mura et al., 2013, Sercombe et al., 2015, Zaleski-Larsen and Fabi, 2016, Daraee et al., 2016, Alavi et al., 2017) to reduce the effective dose.

## Acknowledgments

The authors would like to thank Richardson Patrick Joseph for editorial assistance for his generous support, insightful comments, and suggestions.

## Availability of data and materials

Not applicable

## Authors’ contributions

TPJ designed the study and performed experiments. QZ, WC and SK designed the study and participated in the analysis of results, review and editing of the manuscript. FYK participated in the analysis of results, review and editing of the manuscript. MTZ has contributed in the project administration, interpretation of results and revision of the manuscript. The final manuscript was approved by MH.

## Ethics approval and consent to participate

Not applicable

## Consent for publication

All authors agree to the publication of this article.

